# Is the HIV epidemic over? Investigating population dynamics using Bayesian methodology to estimate epidemiological parameters for a system of stochastic differential equations

**DOI:** 10.1101/219832

**Authors:** Renee Dale, Beibei Guo

## Abstract

Current estimates of the HIV epidemic indicate a decrease in the incidence of the disease in the undiagnosed subpopulation over the past 10 years. However, a lack of access to care has not been considered when modeling the population. Populations at high risk for contracting HIV are twice as likely to lack access to reliable medical care. In this paper, we consider three contributors to the HIV population dynamics: susceptible pool exhaustion, lack of access to care, and usage of anti-retroviral therapy (ART) by diagnosed individuals. We consider the change in the proportion of undiagnosed individuals as the parameter in a simple Markov model. We obtain conservative estimates for the proportional change of the infected subpopulations using hierarchical Bayesian statistics. The estimated proportional change is used to derive epidemic parameter estimates for a system of stochastic differential equations (SDEs). Epidemic parameters are modified to capture the dynamics of each of the three contributors, as well as all their possible combinations. Model fit is quantified to determine the best explanation for the observed dynamics in the infected subpopulations.

**Author summary:** Using a combination of statistics and mathematical modeling, we look at some possible reasons for the reported decrease in the number of undiagnosed people living with HIV. One possibility is that the population of people at significant risk to contract HIV is being depleted (susceptibles). This might happen if significant risk for HIV infection occurs in small percentages of the overall population. Another possibility is that infected individuals lack access to care in some regions due to poverty or other cause. In this case we have to question the accuracy of the estimated size of that population. Finally, most diagnosed individuals report being on medication that reduces their viral load. This greatly reduces their chance to transmit HIV to susceptible individuals. We also combine these possibilities and look at the best explanation for the infected population size.

## Introduction

The human immunodeficiency virus (HIV) progresses in three stages. The first stage lasts approximately 3 months and individuals in this stage are approximately 10 to 25 times more effective at transmitting the disease [16]. The chronic stage can last from 5-10 years without medication [25].This is followed by auto-immune deficiency syndrome (AIDS) and death shortly thereafter [16]. Individuals with HIV may go many years without diagnosis, during which time they may expose uninfected individuals to the disease. Efforts to improve the diagnosis rate include educational programs, as an individual’s perceived risk was shown to be highly correlated with the individual obtaining multiple HIV tests [6, 9, 11, 22]. Several studies have found a 50% reduction in risky behaviors after diagnosis, including safer sex practices, reduction in partner number, and medications that reduce viral load [14, 26, 31]. However, diagnosis events resulting in behavior modification are not thought to be sufficient to eradicate the epidemic [7, 14].

After diagnosis, infected individuals have the opportunity to take anti-retroviral therapy (ART) that reduces their viral load and retards the progression of the disease. The earlier that ART is received the higher the reduction in transmission events. ART therapies could eradicate the epidemic in a population with high prevalence of infection even without the additional effect of behavioral changes [7]. Mathematical models estimate that the HIV epidemic could be reduced to less than 1% of the population infected (elimination phase) by providing ART consistently to newly diagnosed individuals [23]. However, issues with adherence and resistance are well documented in the literature [1–3, 12, 15, 19]. Patients tend to report their adherence as much higher than it actually is, but studies indicate that even low adherence may be sufficient for control of the epidemic [1, 26]. Transmission is rare for individuals on ART, even with relatively high plasma HIV concentrations [17, 20].

The largest barrier to eradication of the epidemic is lack of access to care, including diagnostic services and ART costs or prescriptions. A lack in access to care could create pockets of undiagnosed individuals while the overall trend appears to be a reduction in the size of the epidemic [4]. Various studies report between 50 - 96% of diagnosed individuals in the U.S. rely on public medical care to obtain their ART medications [5, 8, 19, 21, 26]. Access to care remains critical, but this has not been considered when modeling the dynamics of the epidemic [6, 9].

Estimates using CD4 levels of newly diagnosed individuals suggest that the undiagnosed population is decreasing between 2005 - 2013 [24]. CD4 levels can be used to estimate the progression of HIV [27]. We consider three possible causes for this decrease including exhaustion of the susceptible population. The size of the susceptible population is not easy to estimate since it depends on behavior risk. High risk populations include individuals in poverty and men who have sex with men (MSM) [25]. This is particularly critical in the southern U.S., where individuals tend to be poor and lack access to medical care [4, 25]. As the at-risk population decreases the number of diagnoses will also decrease, which will cause the estimated number of undiagnosed individuals to decrease.

An additional possibility is that the reduction in number of diagnoses is due to individuals lacking access to care. HIV is over-represented in impoverished populations where access to diagnosis and treatment may be more difficult to obtain. In this case, the number of newly diagnosed individuals is not representative of the number of undiagnosed individuals, and the estimates will be inaccurate. Finally, the usage of anti-retroviral therapies reduces the viral load and transmission potential of infected individuals.

We use coupled statistical and mathematical methodology to study the relationships between the three hypothesized causes and their respective population dynamics. We use hierarchical Bayesian statistics to get estimates for the size of the infected populations and their proportional changes across the years. These estimates are used to calculate epidemiological parameters for a system of stochastic differential equations. Our motivation to model the system stochastically arises from heterogeneity due to reporting delays associated with population-level data [18]. The resulting simulations give insight into the implications of the estimated undiagnosed population. We hope this study will help inform future efforts to improve the situation of infected individuals and prevent future outbreaks.

## Materials and methods

### Bayesian Statistics

A Markov model where *p_t_* centered at *qp*_*t−*1_ is used to estimate the proportional change in the infected populations over time, where *p_t_* is the proportion in the current year and *q* is the proportional change. These random variables are estimated using Bayesian statistics.

The sampling model is *x_t_ ~ Bin*(*n_t_, p_t_*), where *n_t_* is population size in the current year. The random variable *q* is taken as a hyperparameter for *p_t_*. The random variables to be estimated for each infected subpopulation are *q* and *p_t_*, where t = 2005, …, 2013. We estimate the random variables of undiagnosed and diagnosed subpopulations independently.

### Prior

The prior for the proportional change *q* is a gamma distribution.

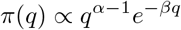

The parameters were chosen so that the prior distribution was centered at the arithmetic estimates of *q* obtained from the CDC [25]. The arithmetic estimates were obtained by calculating:

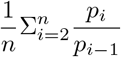

The arithmetic estimate for the undiagnosed *q* (*q_u_*) is 0.979, and for the diagnosed *q* (*q_d_*) 1.025. The priors used were GAM(9.79,10) and GAM(10.25,10) so they were centered at 1.

The prior for the random variable *p_t_*, the undiagnosed proportion, is a beta distribution centered at the previous proportion times *q*. The parameters of the beta distribution are *α* = 0.1*n_t−_*_1_ *× *qp*_t−_*_1_, and *β*= 0.1*n_t−_*_1_ − *α*.

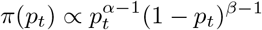

In the case where *t* = 1, the previous undiagnosed proportion is taken to be the expert opinion of 20%, and the diagnosed proportion to be 1 *− p*_0_(*undiagnosed*) [10]. The prior for the diagnosed population is formulated in the same way. Population sizes were considered in units of thousands.

### Likelihood

The likelihood is a binomial distribution, representing the chance of selecting an undiagnosed or diagnosed individual at random from the total infected population. For a given year *t*, the proportion of undiagnosed individuals depends on the total number of individuals:

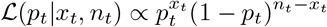
where *x_t_* is the total number of undiagnosed or diagnosed individuals, and *n_t_* is the total number of infected individuals. The likelihood across all the years is the product of each year’s likelihood.

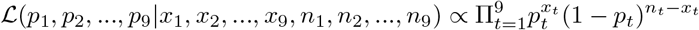

The likelihood for the diagnosed population was formulated in the same way.

### Posterior

The joint posterior distribution is proportional to the priors multiplied by the likelihoods for all 9 years:

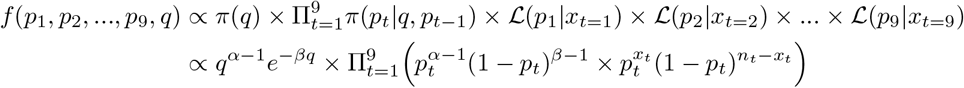

The posterior full conditional of *p_t_* for *t* = 2005, …, 2012 is:

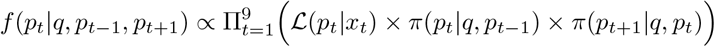

The posterior full conditional of 2013, the 9th year, is:

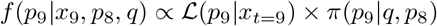

The full conditional of *q* does not have a closed form. The forms of the diagnosed random variables are the same. Random variable estimates were obtained using Metropolis-Hastings nested within a Gibbs sampler over 100,000 iterations with R version 3.3.3 [28]. The proposal distribution was a truncated normal distribution, using package rmutil [29], centered at the previous value of the parameter. Proportions 1 through 9 were sampled consecutively, followed by hyperparameter *q*. The trace plots converged quickly, and the first 2000 samples were removed. Code is available upon request.

### Stochastic Differential Equations

The hyperparameter *q* was estimated to be 0.978 for the undiagnosed population and 1.036 for the diagnosed population. These were used as a boundary to solve for the epidemiological parameters in a simple stochastic differential model.

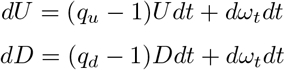

where *U* is the undiagnosed and *D* is the diagnosed populations, and *dω_*t*_ ~ N or*(0*, σ*) is Brownian white noise with units 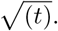 The variance *σ* is chosen to be 10% of the size of the population.

The simplest model is constructed describing the dynamics of the infected subpopulations. The values of parameters transmission (*τ*), diagnosis (*δ*), and death (*ε*) are calculated using the constraining *q*:

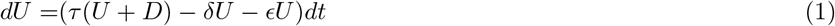

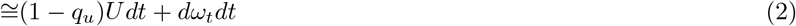

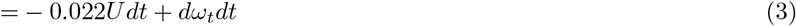

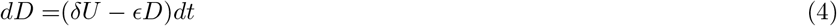

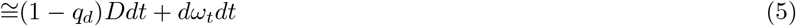

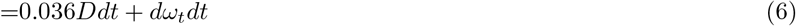

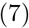

We consider the parameters pseudo-steady state, and use the 2005 population sizes to estimate them. In addition, we assume that the general population are at steady state, and consider only the increased death rate due to infection *ε* as 0.01 [25]. The diagnosis rate is estimated by:

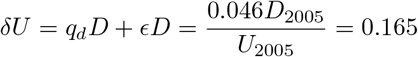

and the transmission rate *τ* is estimated by:

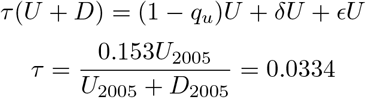

Due to the magnitude of the scale of this system we assume that all events will happen, and the source of the stochasticity is primarily reporting issues. Tau leap algorithm was used to preform the stochastic simulations. A time step of 1 year was selected, and the population at time *t+1* is the numerical solution of the population at time *t* and random noise from a *NOR*(0,*σ*), where *σ* is 10% of the population at time *t*_0_ with units 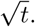 The initial conditions for the infected populations were sampled from the posterior distributions obtained by the Bayesian estimates. Calculations were performed in Matlab [30] and code is available upon request.

The diagnosis rate was calculated using data from [24]. The susceptible population is estimated as twice the national average rate of self-identified MSM among adults. The mortality rate increase due to HIV was estimated using data from [25]. All calculations, including the effective parameter rates, are provided upon request.

## Results

### Bayesian Model

Bayesian estimates for the proportions of diagnosed or undiagnosed individuals was obtained concurrently with the estimated proportional change. The prior distribution was chosen to be a beta for the proportions and a gamma for the proportional changes. The likelihood function was a binomial, representing the chance of randomly selecting a diagnosed or undiagnosed individual from a pool of infected individuals. The posterior did not have a closed form. Due to the symmetry of the posterior samples we summarize them using their mean and variance. The posterior means of the proportions for both undiagnosed and diagnosed estimates are very close to the original data (Fig. ??) [24]. The posterior mean of *q_u_* is 0.96, and *q_d_* is 1.02. This means that 96% of the undiagnosed population is preserved from year to year, or is dropping by about 4% per year. Similarly, the diagnosed population is increasing by 2% per year. Posterior means and variances are given in Table 1.

**Table 1.**
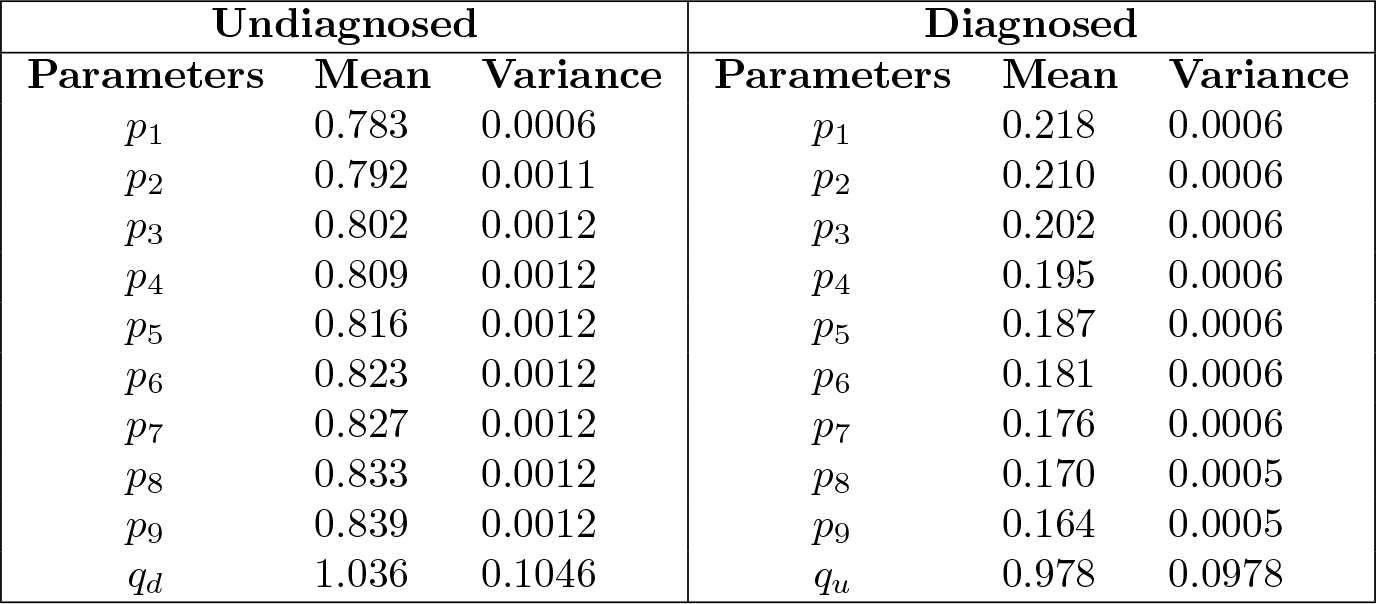
Summary statistics of the posterior distribution.

**Fig 1.**
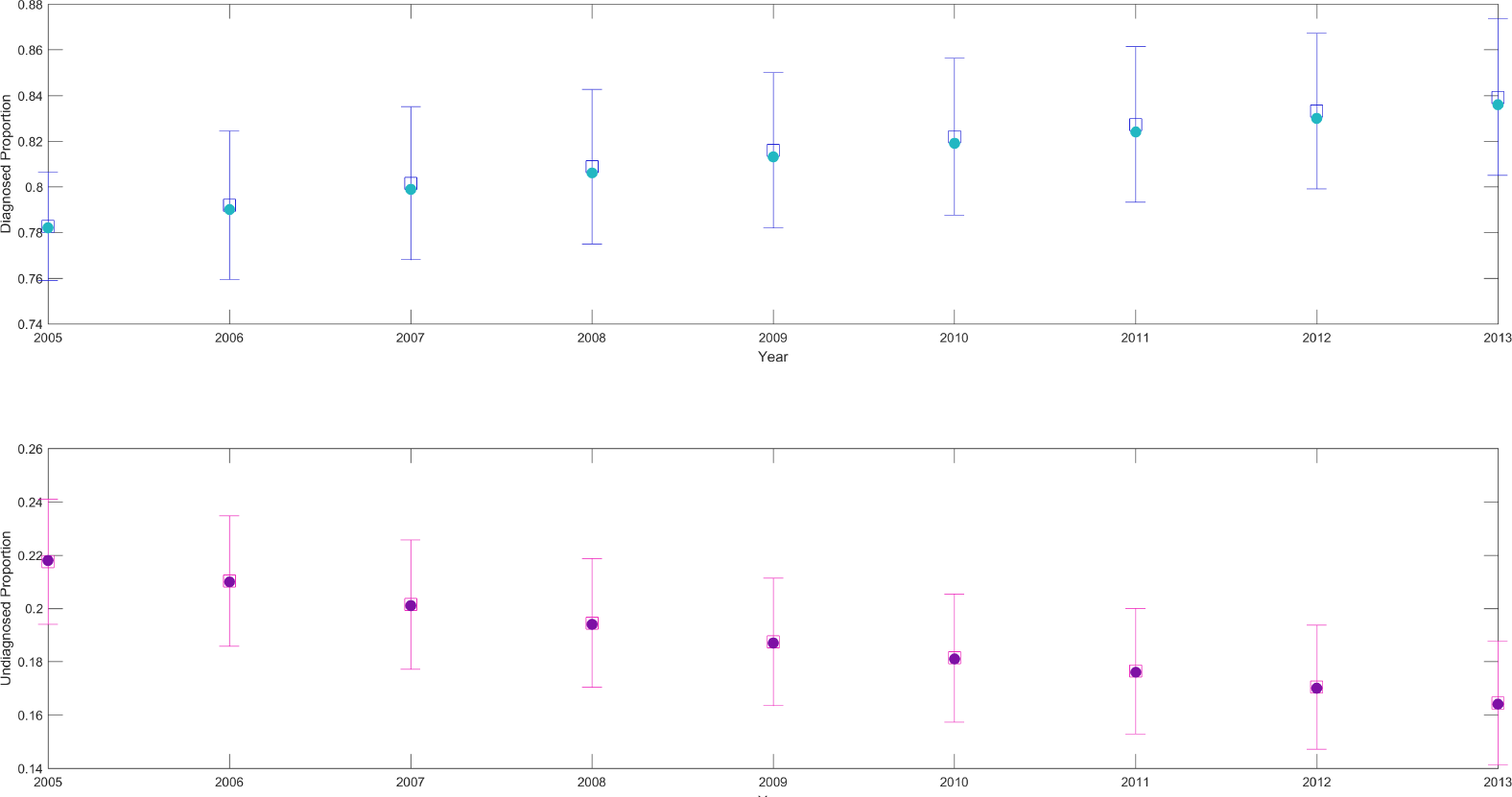
Posterior information obtained from hierarchical Bayesian statistics. Bayesian estimates are shown as hollow squares with error bars showing standard deviations. Estimated proportion of diagnosed (pink) and undiagnosed (blue) populations recover the estimated proportions (solid circles) [24].

### Stochastic Differential Model

The Bayesian estimates of the proportional change in the diagnosed and undiagnosed population from 2005 to 2013 were used to determine the epidemiological parameters for a system of stochastic differential equations. The parameters transmission (*τ*), diagnosis (*δ*), and death (*ε*) were calculated using the proportional changes in the respective population.

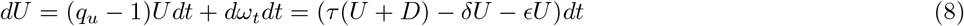

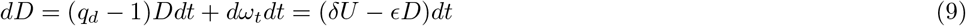

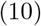

where *U* is the undiagnosed and *D* is the diagnosed populations, and *dω_*t*_ ~ N or*(0*, σ*) is the noise term. These base estimates fit the data very well (Fig. ??).

**Fig 2.**
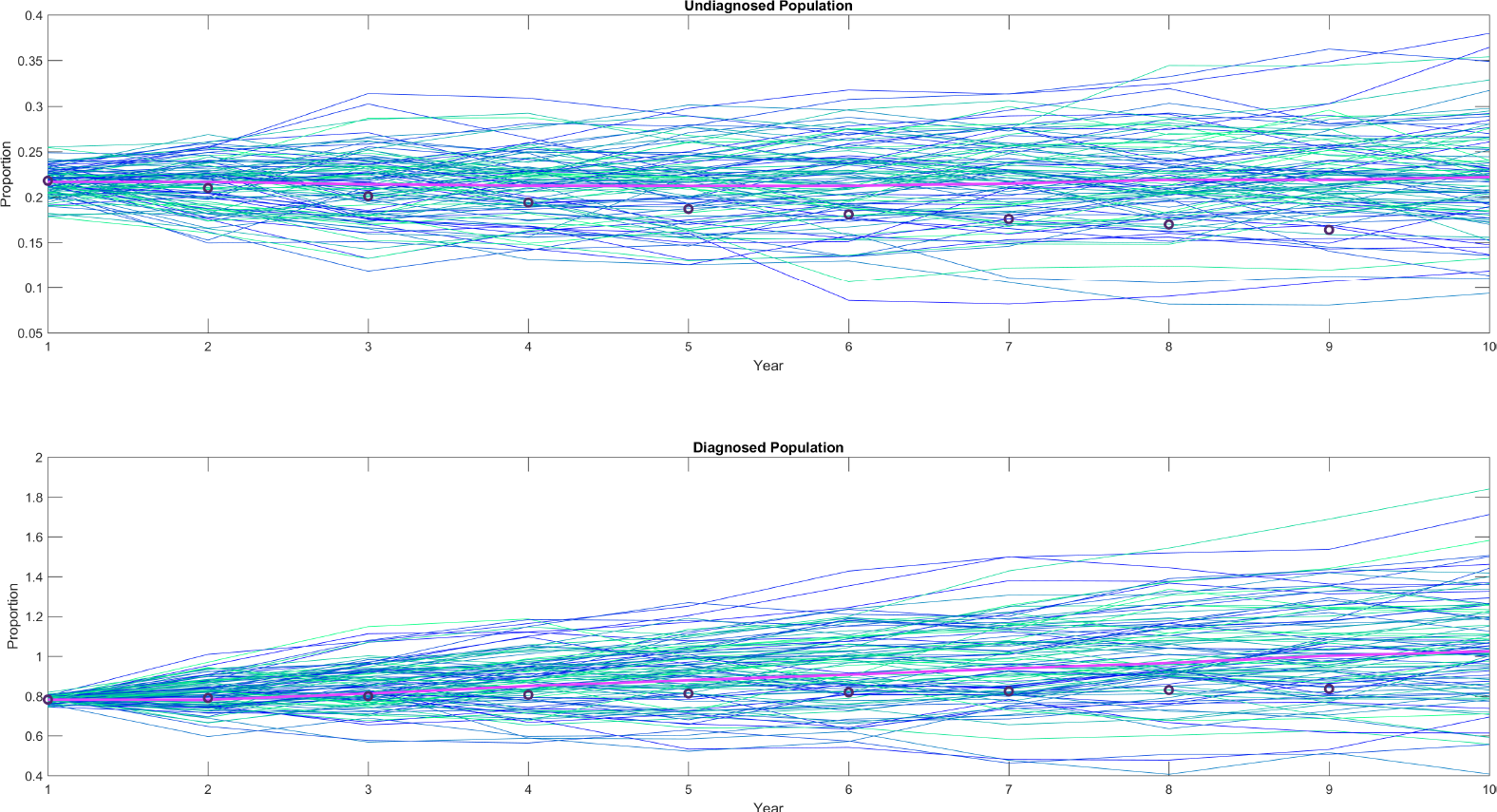
Method Validation. The method was tested by simulating with the epidemiological parameters calculated using the Bayesian estimates of the proportional changes as constraints. The mean of 100 stochastic simulations (pink line) is compared with the data (black circles). Proportions are relative to initial proportion.

### Exhaustion of Susceptibles

In the case where the susceptible population is not much larger than the infected population, the transmission is dependent on the size of both populations. We estimate the susceptible population size as a fraction of the total infected population:

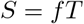

**Table 2.**
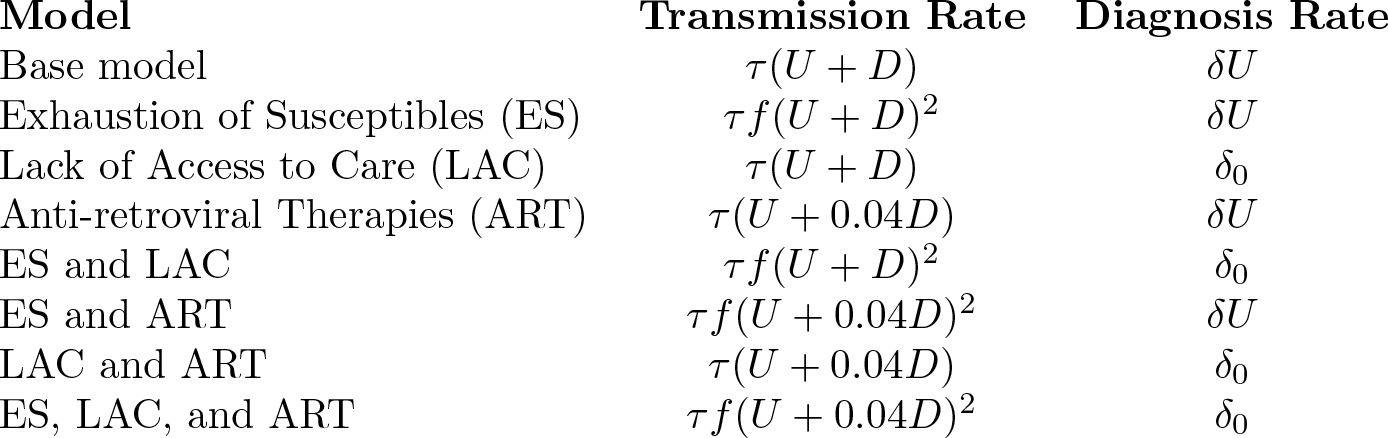
Transmission and diagnosis rates are different under the different hypotheses.

Then this is substituted into the model. The transmission term becomes

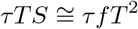

This gives an effective increase of *fτ* in the transmission rate (Table 2). This increase causes the simulations to fail to recover the diagnosed and undiagnosed population dynamics, although the susceptible population does decrease significantly (Fig. ??). This result is intuitive since the infection rate is increased, but the diagnosis rate is not representative of this rate.

**Fig 3.**
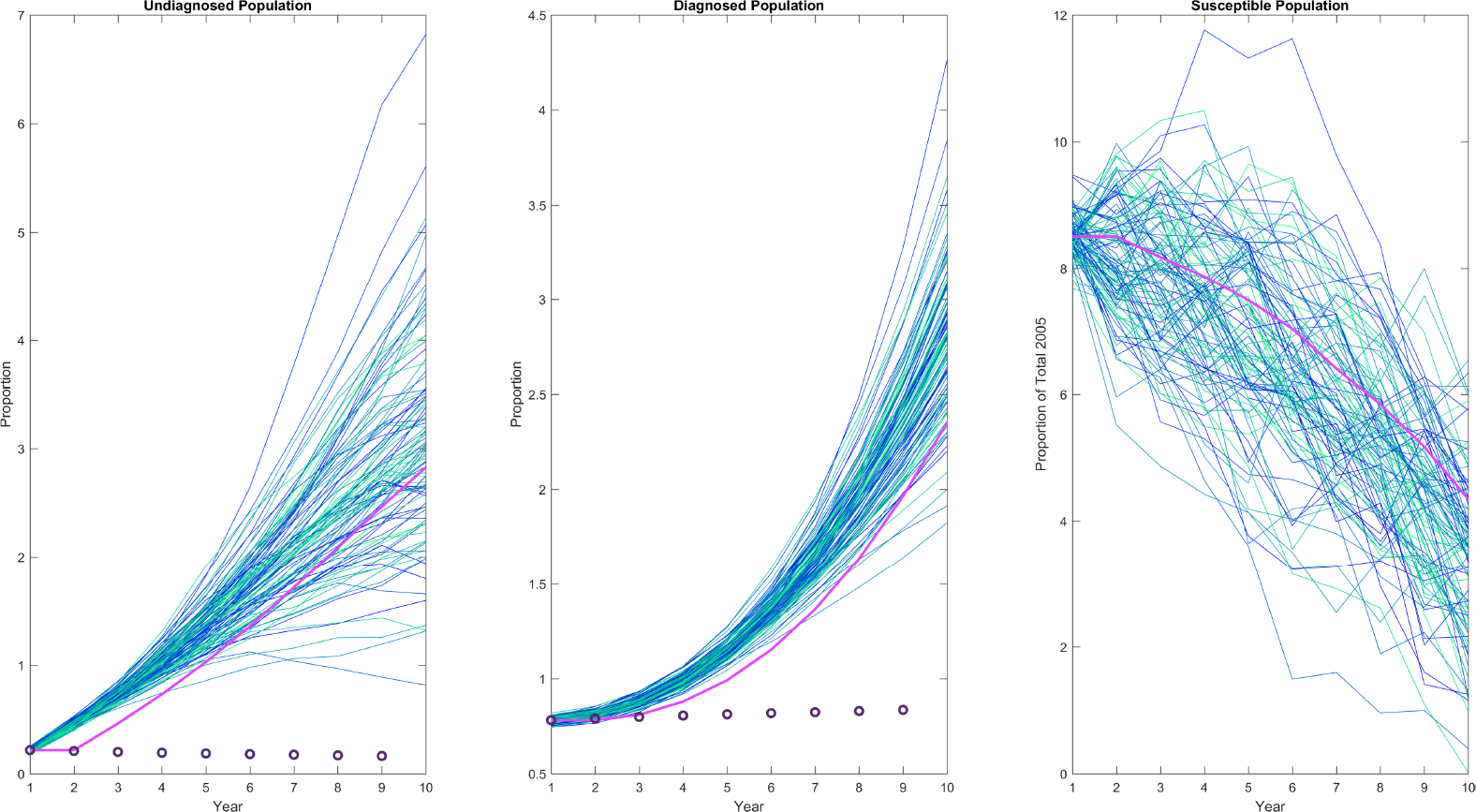
Exhaustion of Susceptibles. Transmission of the disease is altered to reflect the impact of the size of the susceptible population. The mean of 100 stochastic simulations (pink line) is compared to the data (black circles). Proportions are relative to initial proportion.

### Lack of Access to Care

Lack of access to care may be conceptualized as pockets of undiagnosed individuals who are not being diagnosed. To capture this, we consider the diagnosis rate to be independent of the size of the undiagnosed population. The diagnosis rate is estimated as:

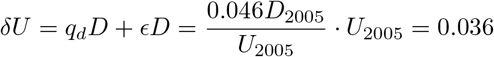

The resulting equation for the undiagnosed subpopulation then becomes:

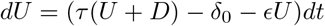

where *δ*_0_ is 0.036. This large reduction in the diagnosis rate recovers the population dynamics well (Fig. 4).

**Fig 4.**
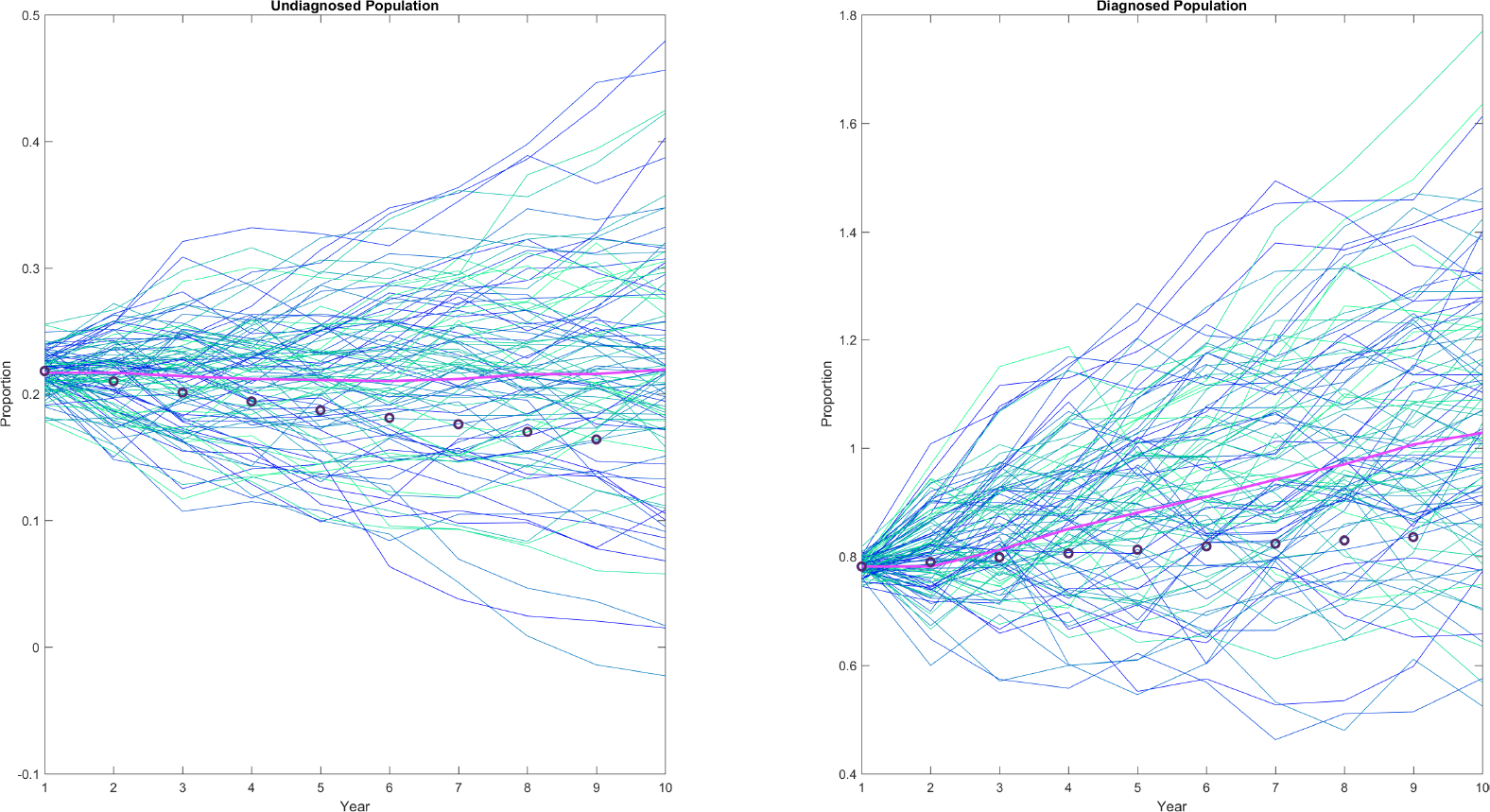
Lack of Access to Care. The effect of undiagnosed individuals lacking access to care affects the rate of diagnosis of the undiagnosed individuals. The diagnosis rate is held constant to reflect this scenario. The mean of 100 stochastic simulations (pink line) is compared with the data (black circles). Proportions are relative to initial proportion.

### ART Usage

Since ART results in a viral load that has low chance of infecting a susceptible individual, we removed these individuals from the pool of infected individuals able to transmit the disease. Since 96% of diagnosed individuals reported taking anti-retroviral therapies in a previous study, the transmission term was modified as follows [26].

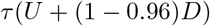

Variable or poor adherence on the part of diagnosed individuals is ignored due to the body of literature indicating that large benefit is gained from even poor adherence [1, 12]. This gives good recovery of both subpopulation dynamics and agrees best with both undiagnosed and diagnosed estimates (Fig. 5).

**Fig 5.**
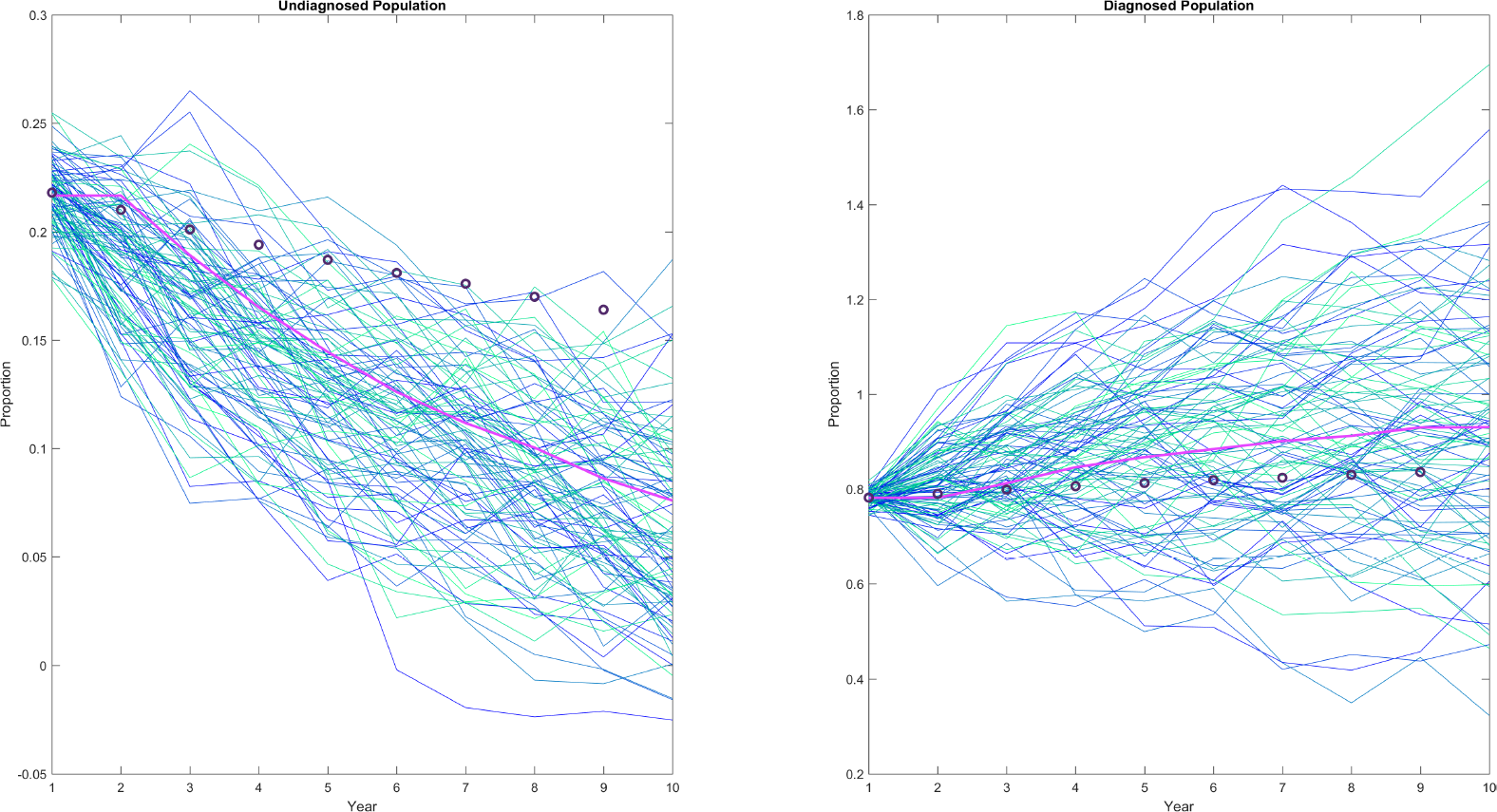
Anti-retroviral Therapy Usage. To reflect the high levels of ART prescription and usage reported by diagnosed individuals this percentage is removed from the pool of diagnosed individuals able to transmit the disease. The mean of 100 stochastic simulations (pink line) is compared to the data (black circles). Proportions are relative to initial proportion.

### Multiple Causes

Since it seems likely that most or all of these scenarios affect the infected population simultaneously, we analyze all their possible combinations (Figure not shown). To determine the best cause, we quantify the goodness of fit by determining the relative likelihood of observing the data given the mean and variance of the stochastic simulations. These probabilities are given in Table 3, as well as the average probability over the 9 years.

**Table 3.**
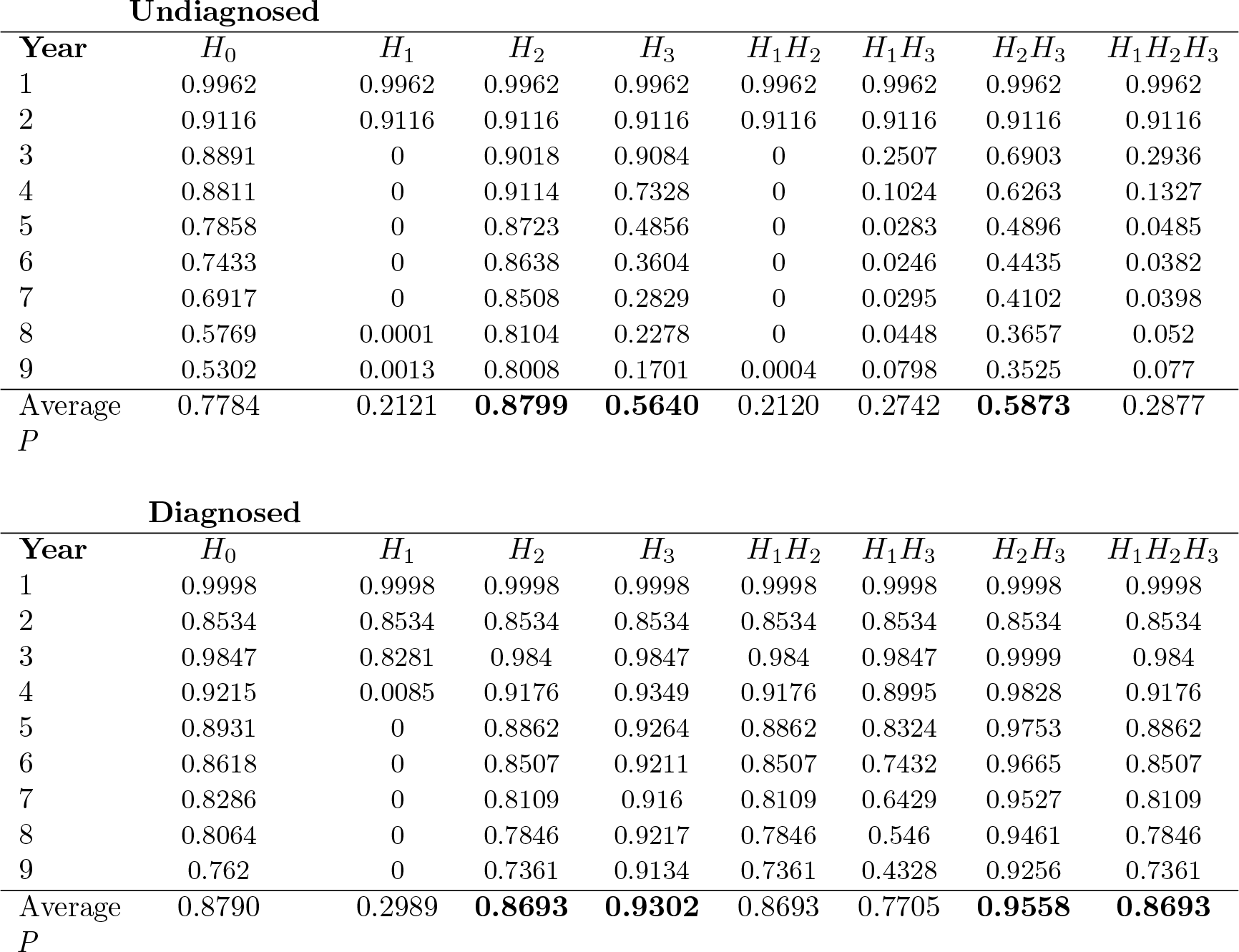
The probability of the data being observed in 100 stochastic simulations compared between the 7 scenarios. The 3 highest average probability over the 9 years is bolded.

## Discussion

We were able to obtain conservative estimates of the proportional changes in the diagnosed and undiagnosed HIV-infected populations using hierarchical Bayesian statistics. Our estimates suggest that the proportion of infected individuals who are undiagnosed is decreasing by approximately 2.2% each year from 2005 to 2013, while the proportion of diagnosed individuals is increasing by approximately 3.6%. We used the proportional change as constraints on a system of stochastic differential equations. This allowed us to estimate the transmission and diagnosis rates. We were able to recover population dynamics using this methodology. To learn more about the cause of the decrease in the undiagnosed population, we considered some scenarios that would affect the epidemiological parameters: exhaustion of the susceptible population, lack of access to care, and reduction in viral load by anti-retroviral therapy.

We were able to recover the diagnosed population dynamics when we altered the parameters to reflect these scenarios with the exception of including exhaustion of susceptibles. Including the size of the susceptible population dramatically increased the transmission rate and caused the size of the infected populations to increase rapidly. In the other scenarios some interesting dynamics could be observed in the undiagnosed population. Lack of access to care was simulated by reducing the diagnosis rate by approximately 13%. This resulted in an improvement in the likelihood of observing the data (Table 3). Anti-retroviral therapy usage also improved the overall recovery, but this effect was weaker for the undiagnosed population dynamics. Although the undiagnosed population size is dependent on the quality of the data available on the diagnosed population of that year, these results indicate that the scenarios that maximizes the probability of observing the diagnosed population also maximizes the probability of observing the diagnosed population estimates.

The observed results suggest that lack of access to care and ART usage contribute to the infected population dynamics. This is not unexpected. Many individuals with HIV are reported to lack access to care [4, 13]. In areas with high poverty rates the death rate of infected individuals is much higher than that of the general population [25]. In 2017 the New York Times reported groups of untreated individuals in the deep south dying due to their lack of access to care [4]. The effect of simulating a lack of access to care suggest this to be a significant contributing factor to the infected population dynamics. Both models and studies have shown that providing ART to infected individuals in the early stages of HIV reduces transmission events and frequency of death due to AIDS [7, 15, 17, 19, 20, 23]. Even poor adherence may be enough to control or eradicate the epidemic and increase quality of life for infected individuals [2, 3, 12]. Greater effort must be made to ensure these populations have access to life-saving treatments.

